# Subcellular localization of glypican-5 associates with dynamic cell motility and cell communication of the human mesenchymal stem cell line U3DT

**DOI:** 10.1101/863720

**Authors:** Masao Takeuchi, Kikuko Takeuchi, Yoko Monobe, Tomoyo Takai, Ritsuko Yamaguchi, Jun K. Takeuchi, Ken-ichi Akagi

**Affiliations:** Section of Laboratory Equipment, National Institute of Biomedical Innovation, Ibaraki-city, Osaka, Japan; Division of Bio-informational Pharmacology, Medical Research Institute, Tokyo Medical and Dental University, Bunkyo, Tokyo, Japan

## Abstract

Glypican-5 (GPC5) is a heparan sulfate proteoglycan (HSPG) localized to the plasma membrane. We previously reported that in the human mesenchymal stem cell line UE6E7T3, GPC5 is overexpressed in association with transformation and promotes cell proliferation by acting as a co-receptor for Sonic hedgehog signaling. In this study, we found using an immunofluorescence microscopy that in transformed cells (U3DT), GPC5 localized not only at primary cilia on the cell surface, but also at the leading edge of migrating cells, at the intercellular bridge and blebs during cytokinesis, and in extracellular vesicles. In each subcellular region, GPC5 colocalized with the small GTPase Rab11. These observations suggest that the colocalization of GPC5 with Rab11 is crucial for its function, and that its activity in each subcellular compartment promotes proliferation of U3DT cells. Our findings indicate that GPC5 plays active and essential roles in regulation of U3DT cell proliferation, and provides several insights into the functions of GPC5 that could be elucidated by future studies.

## Introduction

Glypicans (GPCs) and syndecans (SDCs), which are heparan sulfate proteoglycans (HSPGs) displayed on the surface of most mammalian cells, have long been thought to act as co-receptors for cell-surface receptors in several signaling pathways, including Hedgehog (Hh), Wnt, bone morphogenetic protein (BMP), and fibroblast growth factor (FGF) signaling [1]. The glypican family includes six members (GPC1 to GPC6), each of which is linked to the plasma membrane through a glycosylphosphatidylinositol (GPI) anchor, whereas the syndecan family includes four transmembrane proteins (SDC1 to SDC4) of [2]. Recently, we reported that Glypican-5 (GPC5) is dramatically overexpressed in association with transformation after prolonged culture of the human mesenchymal stem cell line UE6E7T3, and that knockdown of GPC5 expression decreases cell proliferation [3]. GPC5 is overexpressed in rhabdomyosarcomas (RMS), and down-regulation of GPC5 expression by RNAi decreases the proliferation rate of RMS cells [4]. Subsequent work showed that GPC5 stimulates RMS cell proliferation by activating Hh signaling by promoting the binding of the ligand Sonic hedgehog (Shh) to Patched (Ptc), the Hh receptor on the cell surface [5]. Similar evidence has also been obtained in cerebellar granule cell precursors [6] and salivary adenoid carcinoma [7]. Conversely, overexpression of GPC5 inhibits prostate [8] and lung cancer cell proliferation [9]. In non-small cell lung cancer, some reports have suggested that GPC5 is a tumor promoter [10], whereas others insist that it is a tumor suppressor [11,12].

Cell-surface HSPGs also function as potent co-receptors for FGF signaling, as well as Hh signaling; syndecans and glypicans modulate FGF activity by promoting binding of FGF to its receptors (FGFRs) [13]. In particular, they play roles in tumorigenesis and cancer progression. GPC1 is overexpressed in human pancreatic cancer cells [14], breast cancer cells [15], and gliomas [16], and it increases the proliferative response to FGF2, heparin-binding epidermal growth factor-like growth factor (HBEGF), and HGF. In addition, knockdown of HSPGs in these cancer cells decreases the rate of proliferation, suggesting that GPC1 potentiates FGF signaling. Likewise, GPC5 induces a greater increase in the proliferation rate of RMS cell line in the presence of FGF2 [4]. However, GPC5 localizes near primary cilia in RMS [5] and neural precursors [6], indicating that GPC5 interacts with Ptc1 receptor in Hh signaling but not in FGF signaling. We also detected strong staining of GPC5 in the same perinuclear region as concentrated Ptc1 in immunostained U3DT cells [3]. Although these studies clearly establish HSPG as a co-receptor for FGF- or Hh-mediated signaling, the study of SDC4 signaling demonstrates that full activity of FGFs requires not only receptor interaction, but also internalization via HSPG-dependent pathways [17]. In addition, a recent study revealed a role for SDCs in vesicular trafficking and endocytic control on FGF signaling processes [18].

Although we demonstrated that GPC5 participates in U3DT cell proliferation, HSPGs have numerous cellular functions related to modulation of the Hh, Wnt, BMP, and FGF signaling pathways, depending on cell and tissue type. However, most studies of cell-surface HSPGs to date have focused on syndecans, and except for a few examples, less is known about glypicans, particularly GPC5. Therefore, in addition to its role as co-receptor for Hh signaling, GPC5 might have diverse yet heretofore undescribed functions. As a first approach to identifying other GPC5 functions, we investigated the subcellular distributions of GPC5 during various processes in U3DT cells. We reasoned that an understanding of when and where GPC5 localizes within the cell might provide hints about other GPC5 functions.

Here, we show that GPC5 on the cell surface is associated with FGFR1 and small GTPases (Rab11 and ARF6) during cellular migration, and localizes at the midbody area and plasma membrane blebs during late cytokinesis. In addition, we found many vesicles containing GPC5 in U3DT cell-culture medium. Together, these results have implications regarding the recycling endosomal dynamics of GPC5 in U3DT cells.

## Materials and methods

### Cell culture

The human mesenchymal stem cell line UE6E7T-3 (JCRB1136) which is the same to U3-A in Fig. 1 and their transformed derivative U3DT (JCRB1136.01) were cultured in DMEM containing 10% FBS as described in a previous report [3].

### Preparation of extracellular vesicles (EVs)

EVs were prepared from conditioned medium of U3DT cells cultured for 2 days in DMEM with 0.1% FBS, which had previously been centrifuged at 105,000 *g* for 90 min. The conditioned medium was centrifuged at 800 *g* for 30 min, and the supernatant was filtered with an 0.45 μm Millipore filter and centrifuged at 17,000 *g* for 90 min [19]. The pellet was suspended in PBS or appropriate culture medium and used for immunofluorescence or incorporation assays in UE6E7T3 cells. One milliliter of EV was prepared from 100 ml of conditioned medium of U3DT, and the concentration of the resultant EV solution was 0.1 µg/ml, as determined using a NanoDrop 2000 (Thermo Fisher Scientific, Japan) using BSA as the standard solution. UE6E7T3 cells were cultured with 0.3 µg EVs in 2 ml DMEM. After 1 day, the cultured cells were characterized by immunofluorescence staining.

### Immunocytochemistry

Cells cultured on coverslips were fixed in 4% paraformaldehyde in PBS washed in phosphate-buffered saline (PBS), washed with PBS and then blocked with 1% BSA in PBS. For detection of Rab11, furthermore, samples were fixed with 10% TCA and washed with 50 mM NH_4_Cl and permeabilized with 0.05% Triton-X100 [20]. The cells and extracellular vesicle (EV) were subjected to indirect immunofluorescence staining. Antibodies and fluorescence-conjugated reagent used in this study are as follows; Anti-GPC5 antibody (MAB2607, R&D Systems, Inc.), Alexa Fluor 594-conjucated anti-GPC5 antibody (R&D Systems, Inc.), Alexa Fluor 647-conjucated anti-GPC5 antibody (R&D Systems, Inc.),

Anti-FGFR1 Xp Rabbit mAb (D8E4, Cell Signaling Tec.), Anti-Rab 11A antibody (A-6: sc-166912, Santa Cruz Biotech), Anti-acetyl-alpha-tubulin antibody (D20G3, #5335, Cell Signaling Tec.) rabbit mAb, Alexa Fluor 488-conjucated Wheat germ agglutinin (W1126: Invitrogen), Anti-CD63 (MX-49.129.5: sc-5275, Santa Cruz Biotech), Anti-CD63 (cl 3-13: Fuji Film Co.), Anti-ARF6 (3A-1: sc-7971, Santa Cruz Biotech,), Anti-FGFR-1 (D8E4) XP Rabbit mAb (Cell Signaling). Alexa Fluor 488 conjugated-anti-mouse IgG(H+L), F(ab’)2 fragment (#4408, Cell Signaling), Alexa Fluor 488 conjugated-anti-rabbit IgG(H+L), F(ab’)2 fragment (#4412, Cell Signaling), Alexa Fluor 594 labeled goat anti-mouse IgG IgG(H+L), F(ab’)2 Fragment (A-11020, (Molecular Probes, Inc.), Alexa Fluor 647 conjugate-Anti-Rabbit IgG(H+L), F(ab’)2 Fragment (#4414, Cell Signaling). All antibodies were used at a dilution of 1:100 from 0.2 to 1 μg/ml stock. Immunostained cells or EVs were mounted using ProLong Diamond Antifade Mountant with DAPI (Invitrogen) and visualized on a confocal fluorescence microscope.

### Confocal microscopy

An inverted laser scanning microscope (TCS SP8; Leica, Germany), equipped with a 63x oil objective (NA = 1.4) was used to visualize cells or EVs. For quantitative analysis of fluorescence intensity, fluorescence images were obtained with an extremely light-sensitive HyD detector: intensities at 488, 568, and 594 nm were collected in photo-counting mode. Images were 1024 × 1024 pixels and were collected as Z-stacks (Z-step size, 0.24 μm; zoom, 1.5–10). The sum of fluorescence intensity was calculated from the optimum intensity of z-stack images of each cell, which was expressed as the pixel sum of each cell, using the LASX software (v.3.4, 18368.2) [21]. For quantitative detection, samples were stained under the same conditions, and immunofluorescence images were obtained under the same light and detector conditions on the same day. The background pixel sum of each cell was below 5%.

### Flow cytometry

UE6E7T-3 or U3DT cells treated with or without siRNA-RAB11A for 3 days were harvested with trypsin and were fixed with 4% paraformaldehyde-PBS. The permeabilized cells were suspended in 0.1% BSA-PBS at a concentration of 5– 10 × 10^5^ cells/ml. One hundred microliters of each cell suspension was mixed with 5 µl antibody diluted 20-fold in PBS containing 5% FBS (5% FBS-PBS). After incubation overnight at 4°C, the cell suspension was washed twice with 5% FBS-PBS and shaken with secondary antibodies for 4 h at 4°C.

Data acquisition from 10,000 cells per sample was performed on a FACSAria (BD Biosciences), and the resultant data were analyzed using the FlowJo software (TOMY Digital Biology). The antibodies used in this test were as follows: anti-GPC5 mouse antibody, anti-Rab11A rabbit antibody, Alexa Fluor 488-conjugated anti-mouse antibody, and Alexa Fluor 594-conjugated anti-rabbit antibody.

### siRNA treatment

U3-DT cells were seeded in 6-well chamber slides (5 × 10^3^ cells per well) and cultured in DMEM containing 10% FBS. The following day, cells were transfected with 100 or 200 nM Accell Human Rab11A siRNA SMARTPool (Thermo Fisher Scientific) with 0.1 % FBS, as described previously [3]. After 72 h, cells in each well were fixed with 4% paraformaldehyde and characterized by immunofluorescence.

### Transmission electron microscopy

A 5 μl aliquot of an EV sample was applied to a glow-discharged carbon film grid, stained with 1% (w/v) uranyl acetate solution, and rinsed with distilled water. Grids were examined using a Hitachi H-7650 electron microscope with an acceleration voltage of 80 kV. EV diameters were calculated from ten representative images using the Hitachi EM viewer Ver 03.01 software.

### Statistical analysis

Data were represented as means ± S.D. Differences between means of individual groups were assessed by Student’s two-tailed unpaired t-test. p < 0.05 was considered statistically significant.

## Results

### GPC5 is dramatically overexpressed in association with transformation of UE6E7T-3 cells

In a previous study, we performed whole-transcriptome analysis of the human mesenchymal stem cell line, UE6E7T-3, over the course of transformation. The results revealed that GPC5 was dramatically overexpressed in association with transformation [3]. Fig. 1 shows glypican expression at four stages of culture spanning 295 population doubling levels (PDL). GPC5 was expressed at low levels at the early stage (PDL 60–90, sample U3-A) and overexpressed at the late stage (PDL 231–295, sample U3DT) of long-term culture with increasing the proliferation from the previous report [3]. Other glypicans (GPC1 to GPC4 and GPC6) were expressed at low levels throughout the culture period.

**Figure 1.**
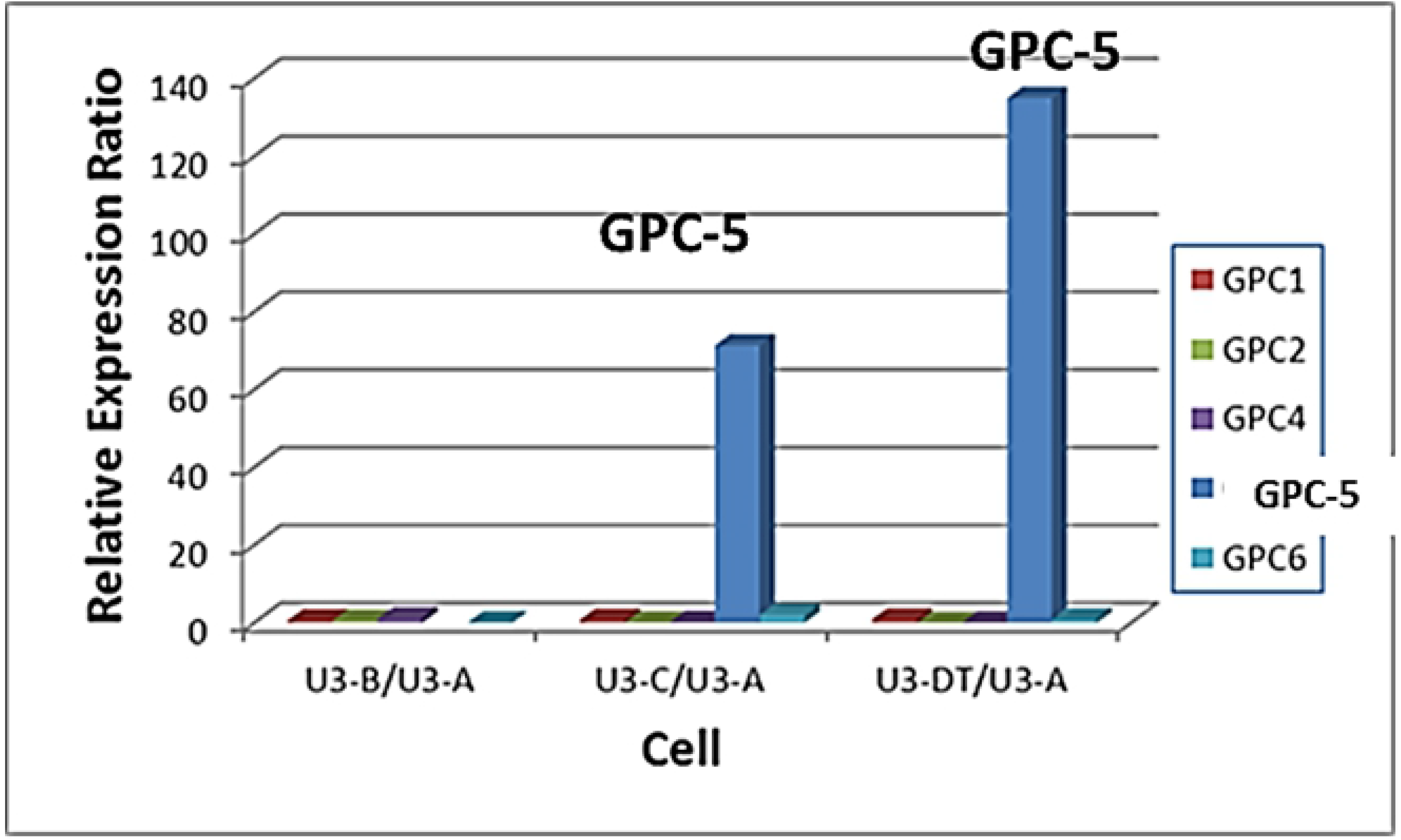
Alterations in GPC5 expression during long-term culture. Gene expression of glypicans (GPC1 to GPC6) at four culture stages were analyzed using data in the DDBJ database of the National Institute of Genetics (http://www.ddbj.nig.ac.jp/), accession number: DRA000533 [3]. Each value is shown relative to the corresponding value in U3-A cells (culture stage 1). GPC1, red; GPC2, green; GPC3, no expression; GPC4, purple; GPC5, blue; GPC6, light blue. As shown in the previous report [3], PDL of U3-A, U3-B, U3-C and U3DT cells are 60-90, 91-150, 151-230 and 231-295, respectively.

### Cell-surface GPC5 colocalizes with FGFR1, Rab11, and ARF6

The FGF–FGFR signaling pathway is involved in cell migration during wound healing [22], and chondroitin and dermatan sulfate or SDC4 modulate FGF-induced cell migration [23,24]. To investigate the interaction between GPC5 and FGFR1, we used immunofluorescence microscopy to examine the localization of endogenously expressed GPC5 and FGFR1 in U3DT cells under steady-state culture conditions. In interphase cells, strong staining patterns of colocalization of GPC5 and FGFR1 were observed in the perinuclear region, and a punctate vesicular pattern was dispersed throughout cell surface. Puncta were detected at the leading edges, such as the tip at the front of an elongating cell (Fig. 2, top panel), blunt-ended protrusions at the front of lamellipodia (Fig. 2, second panel), and region adhered to substratum of a glass coverslip when both daughter cells separate during cytokinesis, as if they might aid in the mechanical separation of the two daughter cells (Fig. 2, third panel). These patterns suggest that GPC5 modulates FGF-mediated cell migration.

In addition, Rab11 and ARF6 GTPases have been implicated in regulation of cell motility [25]. In particular, they regulate the recycling of plasma membrane receptors. Hence, we tested whether GPC5 also interacts with ARF6 or Rab11 in U3DT cell migration. As shown in Fig. 2 (fourth and two bottom panels), both ARF6 and Rab11 were also colocalized with GPC5 at the leading edge of migrating U3DT cells. Based on the results, it is possible that GPC5 and FGFR1 are transported to the plasma membrane by the endocytic recycling pathway and may promote FGF-mediated cell migration.

**Figure 2.**
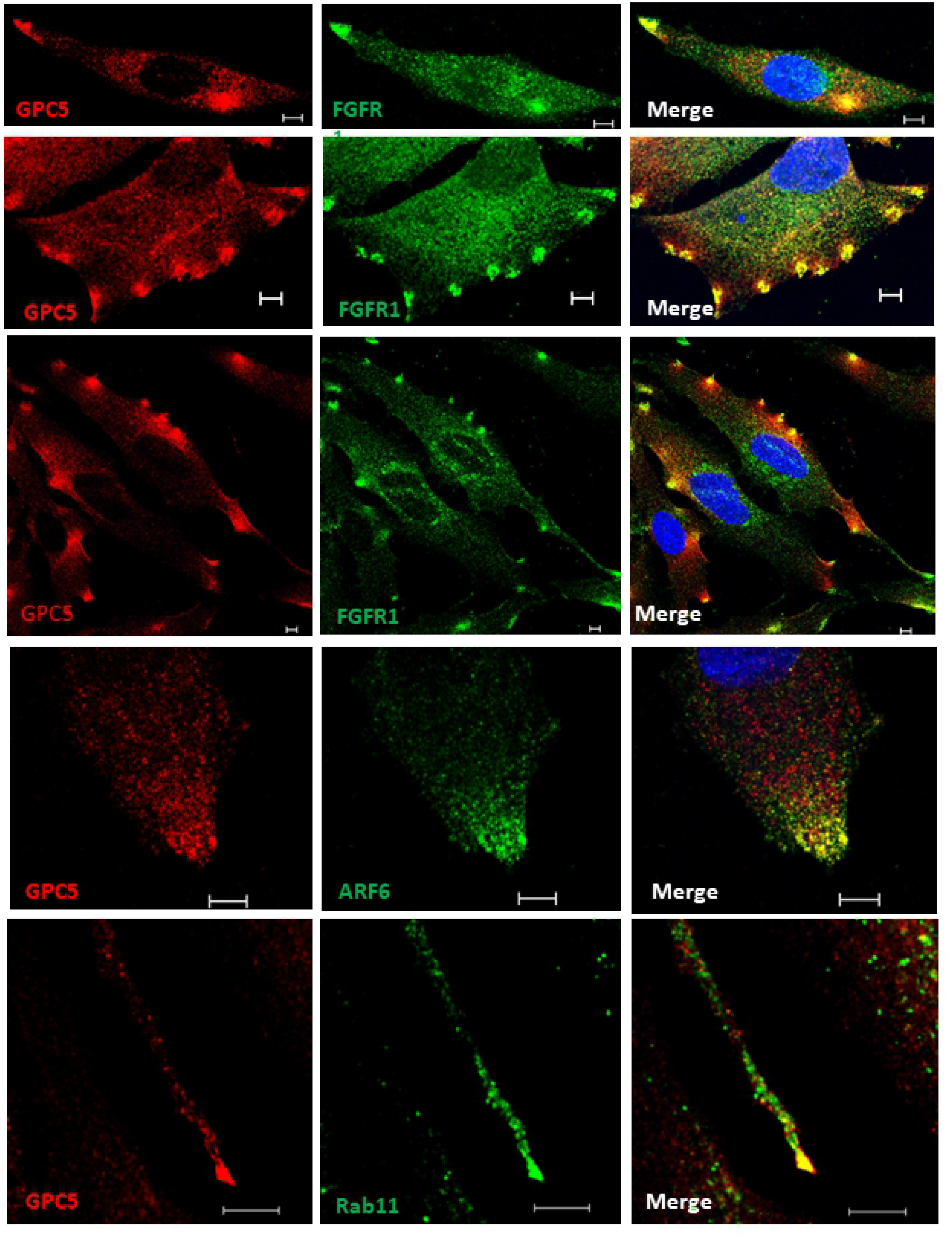
GPC5s localize to the leading edges of migrating cells along with FGF receptor1, Rab11, or ARF6. Double immunofluorescence labeling of U3DT cells with anti-GPC5 (red) and anti-FGFR (green) (top three lines), with anti-GPC5 (red) and anti-ARF (green) (fourth line), or with anti-GPC5 (red) and anti-Ra11A (green) (bottom line) antibodies. Positive spots (yellow) in each merged image were clearly visible in U3DT cells. Scale bar, 5 μm.

### Subcellular localizations of GPC5 during cell division

To test the interaction between GPC5 and FGFR1 during cell division, we first examined the distributions of endogenous membrane GPC5 and FGFR1 in non-permeable U3DT cells by immunostaining. Double staining for GPC5 and FGFR1 revealed that in interphase cells, the two proteins were colocalized at perinuclear regions and leading edges (Fig. 3A). By contrast, the staining patterns in mitotic cells were quite distinct from those in interphase cells. In rounded metaphase cells, strong staining was observed in one or two regions (Fig. 3A, second line); during mitosis, staining was detected adjacent to the cleavage furrow (Fig. 3A, third line). In later stages of cytokinesis, strong staining for both GPC5 and FGFR1 accumulated at the intercellular bridge. At the last step of cytokinesis (abscission), strong GPC5 was also detected in bleb protrusions of both daughter cells (Fig. 3A, abscission). On the other hand, in the cells permeabilized after fixation (Fig. 3B), both GPC5 and FGFR1 exhibited only punctuated staining patterns dispersed throughout interphase cells. At anaphase, however, GPC5 accumulated at the equatorial plane of the cell cortex with FGFR1, and at telophase, GPC5 staining became distinctly concentrated around the central spindle and intercellular bridge (Fig. 3B, third line), whereas FGFR1 staining dispersed into puncta throughout the cell. Similarly, in the last stages of cytokinesis, GPC5 accumulated at high levels accumulated in membrane surface blebs, whereas FGFR1 did not. These results indicate that GPC5 may be transported and play significant roles, some of which involve an interaction with FGFR1, at both the midbody and abscission.

**Figure 3.**
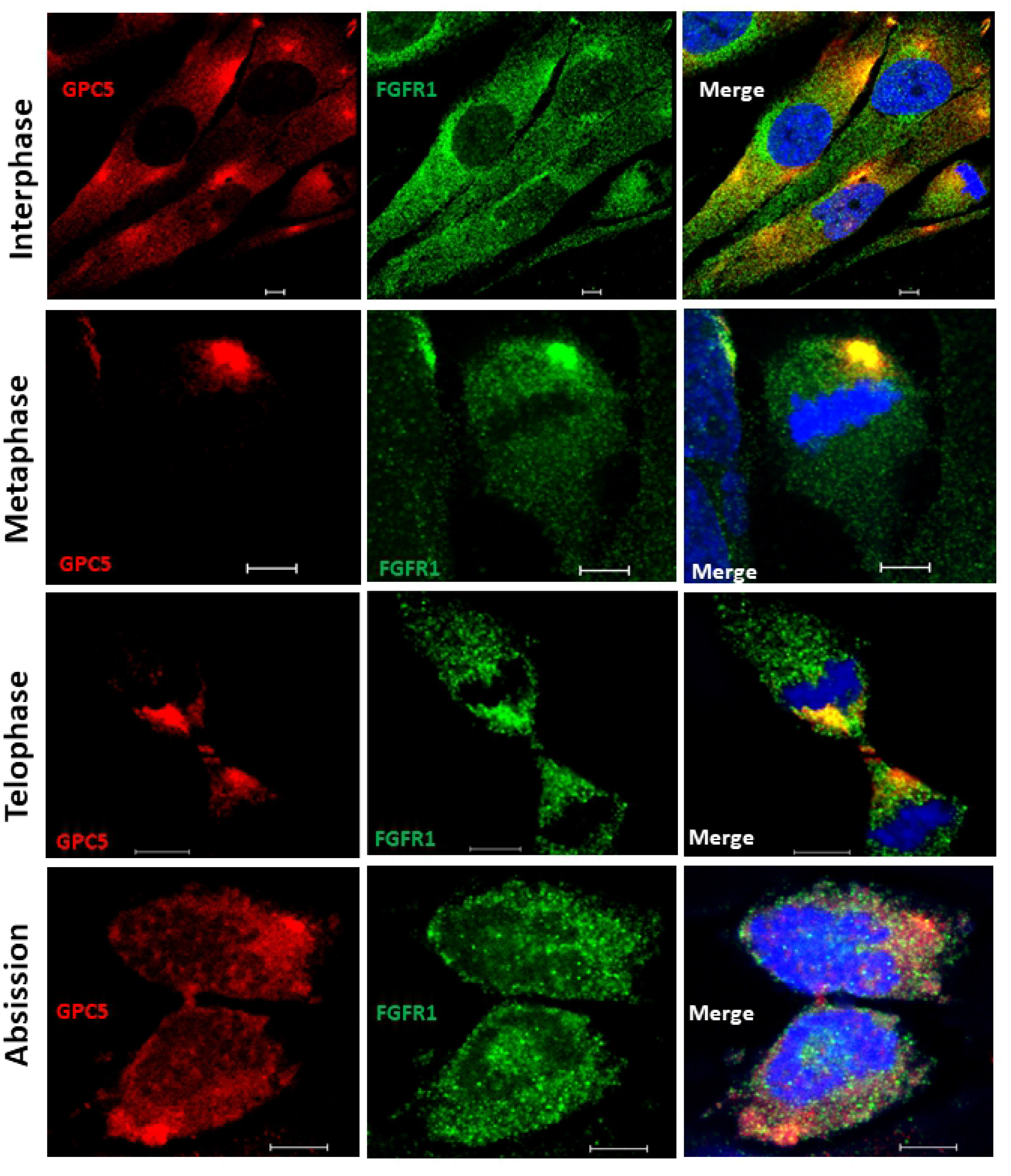
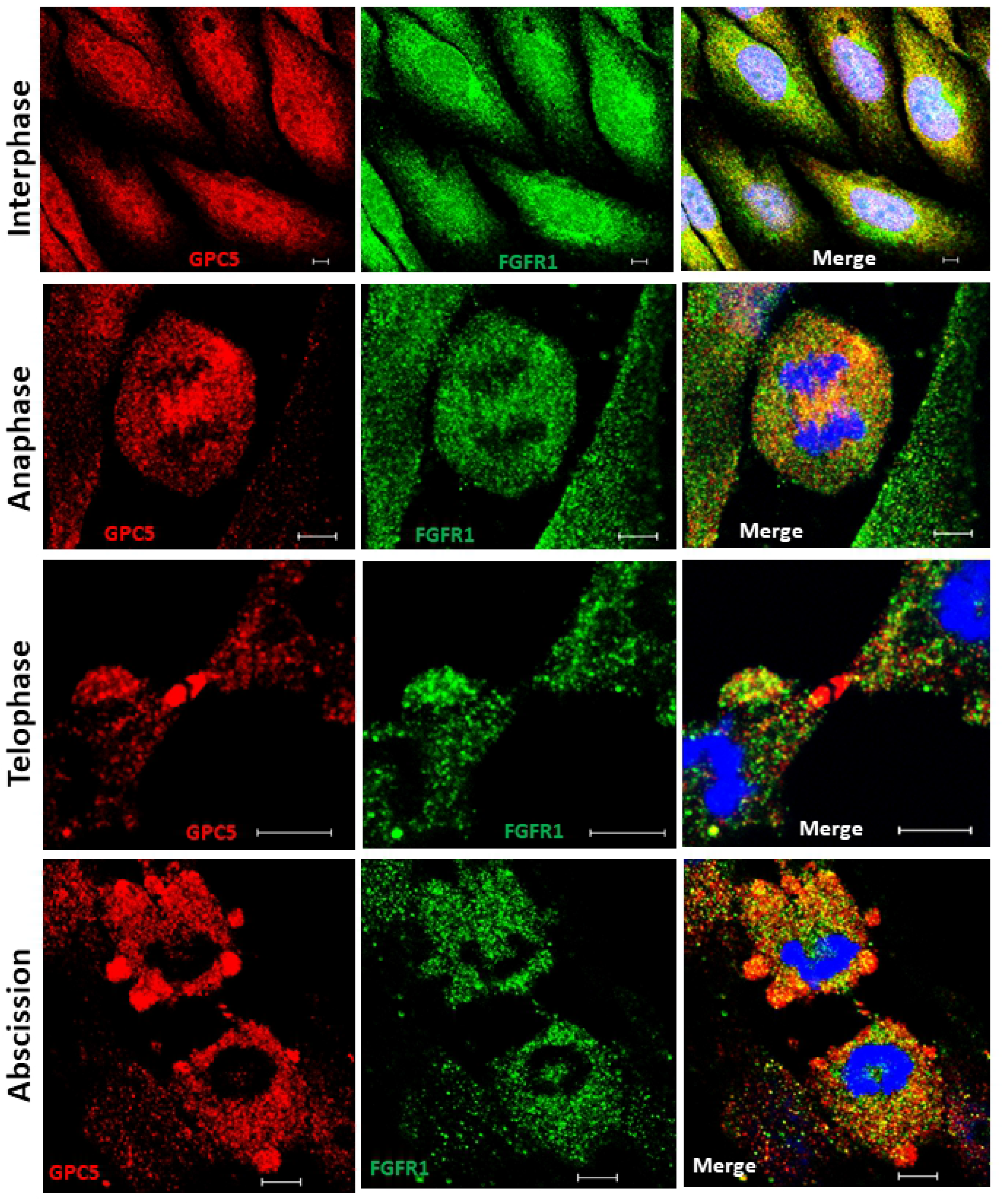
Dynamic localization of GPC5 during mitosis and cytokinesis. (A) U3DT cells at different stages of mitosis and cytokinesis were fixed but not permeabilized, stained with anti-GPC5 (red) and anti-FGFR1 (green) antibodies, and counterstained with DAPI stain (blue). Yellow represents the degree of colocalization. Scale bar, 5 μm. (B) Cells expressing the same marker as in (A) but with permeabilized cells. Scale bar, 5 μm.

### Midbody localization of GPC5

The considerable accumulation of GPC5 at the midbody during telophase was very interesting to us. Large numbers of components are localized within the midzone during telophase [26]. Among them, Rab11, one of the best-studied Rab small G proteins, associates with recycling endosomes and traffics into the midbody during cytokinesis [27,28]. To explore the significance of GPC5 localization at the midbody, we examined the colocalization of GPC5 with tubulin or Rab11. Two faint bands outside the dark zone of microtubules, where tubulin staining did not appear, represented Rab11. By contrast, Rab11 was highly enriched within punctate structures in the proximity of the cleavage furrow, where FGFR1 was also present, although it appeared not to associate with Rab11 (Fig. 4A). To further confirm this observation, we used WGA (fluorescent wheat germ agglutinin [29], a membrane marker that distinguishes cell-surface membrane from cytosolic components. Rab11 staining could also be observed throughout the intercellular bridge but not at the center of the bulge (dark zone) (Fig. 4B). Interestingly, GPC5 also localized there in a pattern that partially overlapped with that of Rab11 (Fig. 4C). The spatial distributions of GPC5, Rab11, and microtubules are shown in Fig. 4D, which confirms the partial overlap of GPC5 with Rab11. These results demonstrate that both GPC5 and Rab11 localize to the intercellular bridge during telophase, but their interaction remains obscure.

**Figure 4.**
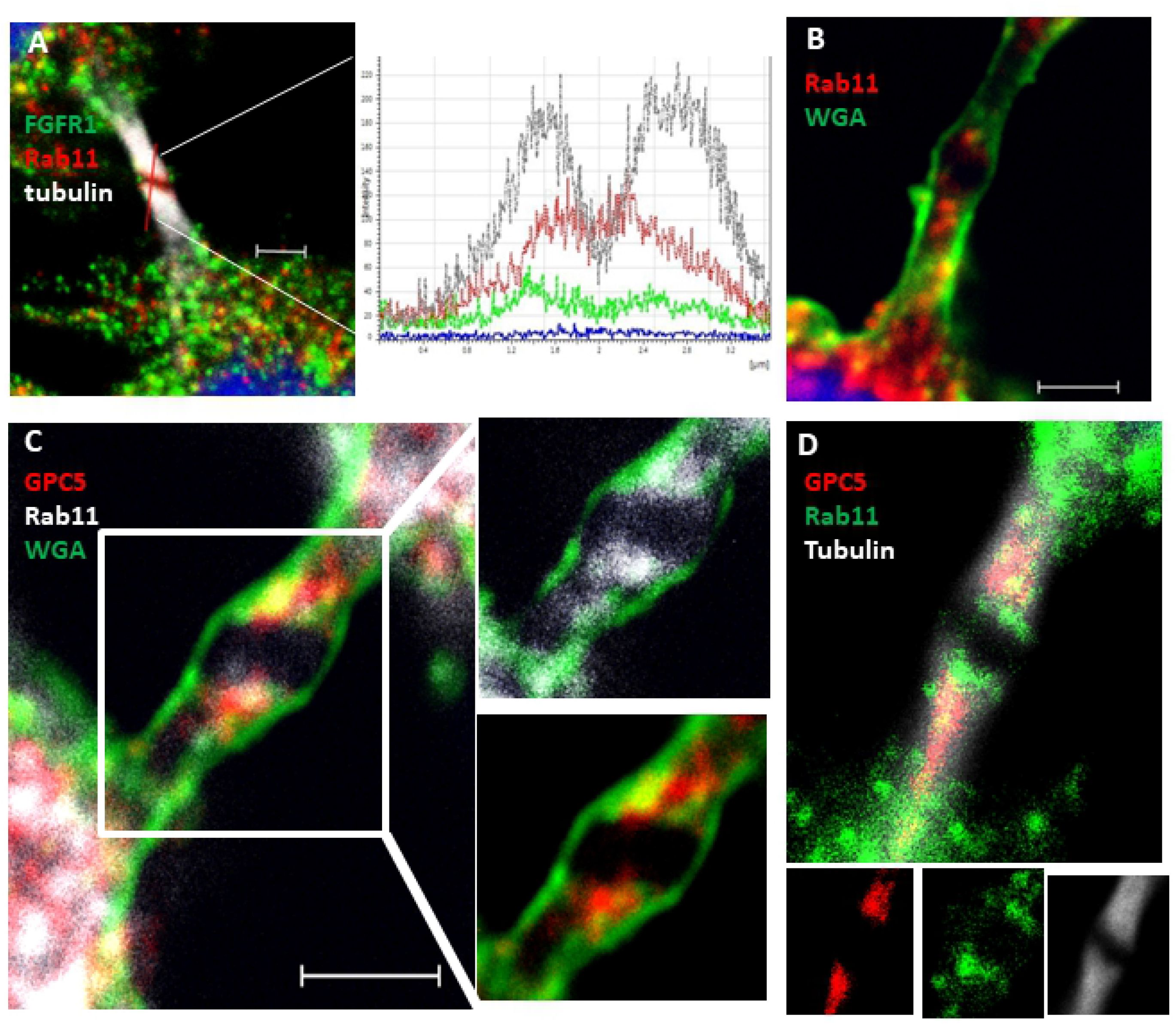
GPC5 localizes to the intercellular bridge. (A) Rab11 localized on both sides (red) of the midbody dark zone and partially overlapped with microtubules (grey). Localizations of proteins on midbody microtubules were determined and compared by line scans. Microtubules and Rab11 peaked at the same positions, where the microtubule signal was high and the FGFR signal was low. (B) Rab11 localization (red) adjacent to the midbody. Plasma membrane was stained with WGA (green). (C) GPC5 (red) colocalized with Rab11 (grey) at the midbody. (D) GPC5 (red) colocalized with Rab11 (green) on midbody microtubules (grey). Scale bar, 2 μm.

### Bleb dynamics in cytokinesis

Membrane blebs have been observed at the poles and midbody of dividing cells [30]. In the present study, U3DT cells spontaneously exhibited dynamic membrane blebs during later stages of mitosis (Fig. 3A–B). Figs. 5A–D show WGA fluorescence staining of U3DT cells during mitosis; membrane blebbing occurred over the entire surface at telophase, similar to exocytic bursts of membrane (Fig. 5D). In some blebs, GPC5 was strongly fluorescently labeled (Fig. 5E), whereas FGFR1 or Rab11 staining was of similar intensity in blebs and in the cytosol inside the surface membrane (Fig. 5F and 5G).

**Figure 5.**
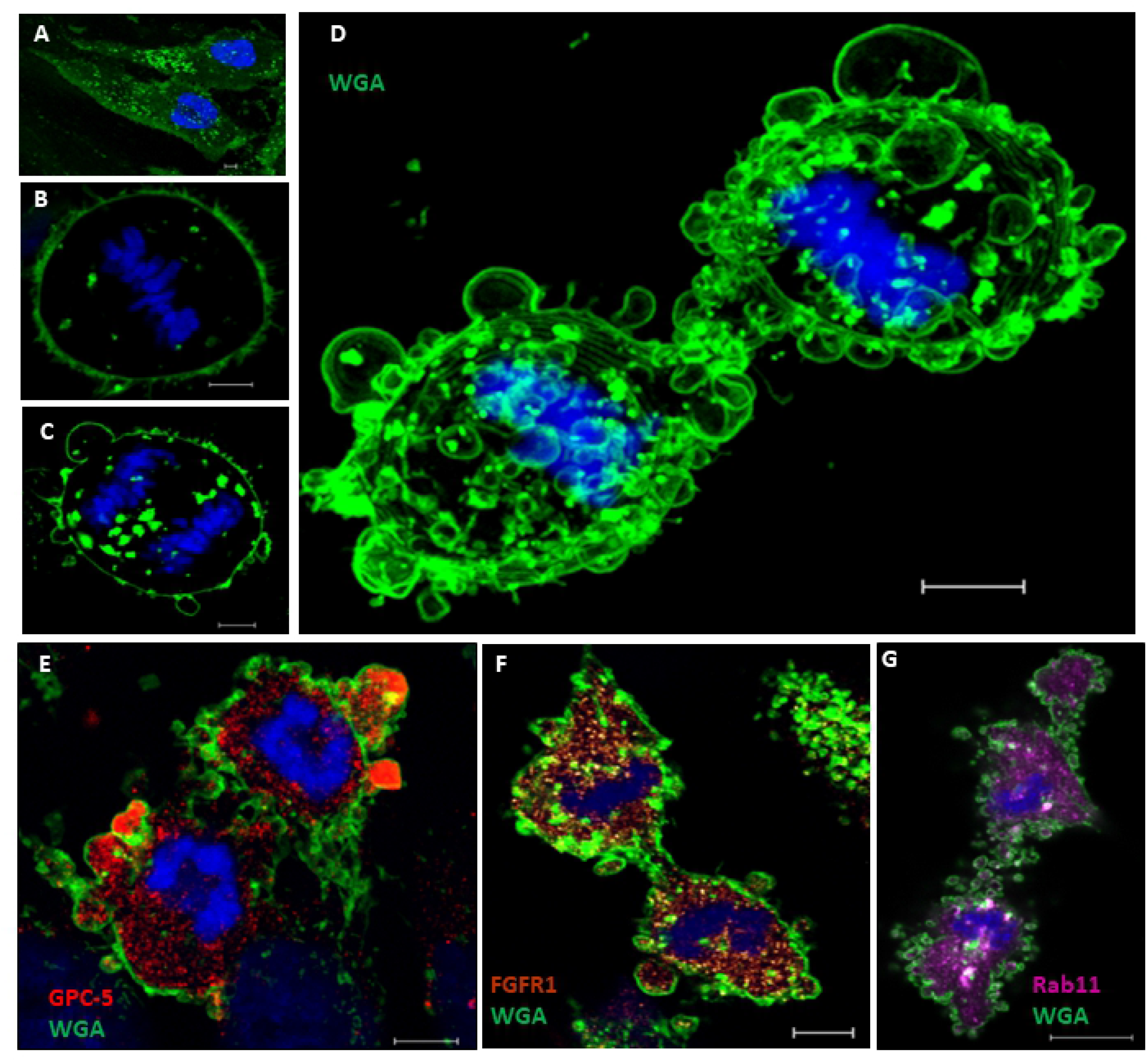
Localization of GPC5 and other markers in blebs of U3DT cells at telophase. (A–D) Images of staining with Alexa Fluor 488-conjugated WGA and DAPI at interphase (A), metaphase (B), and anaphase and telophase (C–D). (D) Maximum projection of 15 Z-staged-images stained with Alexa Fluor 488-WGA and DAPI. (E) U3DT cell at telophase, stained with anti-GPC5 antibody (red) and Alexa Fluor 488-WGA (green). (F) Blebs of U3DT cell at telophase stained with anti-FGFR1 (brown), Alexa Fluor 488-WGA (green), and DAPI (blue). (G) Blebs of U3DT cells at telophase stained with anti-Rab11A antibodies (magenta), Alexa Fluor 488-WGA (green), and DAPI (blue). Scale bar, 5 μm.

The role of blebbing in cytokinesis remains obscure, but it has been speculated that blebbing might contribute to separation of daughter cells via the vigorous motile activity induced by actomyosin contractile system and membrane reservoirs (S1 Fig.). Blebbing appears to correlate with membrane transport. Therefore, we investigated whether the recycling endosome marker Rab11 is present in blebs during cytokinesis. Rab11 was uniformly distributed in blebs as well as cytoplasm (Fig. 5G). Double labeling of GPC5 and Rab11 revealed strong punctate GPC5 labeling at the leading edges of blebs, whereas Rab11 was found partial distribution in blebs, suggesting that the two proteins partially overlap in blebs (S2 Fig.). To further confirm this, we examined the effects of Rab11 depletion on GPC5 localization and bleb formation during cytokinesis. Immunostaining revealed a significant reduction in Rab11 intensity (Fig. 6B–C, 20% decrease), whereas a considerable increase in GPC5 intensity was detected following treatment with RAB11A-siRNA (Fig. 6E–F, 200% increase). Double labeling of cells in suspension (trypsin treatment) with GPC5 and Rab11 yielded results similar to those obtained in adhered cells (Fig. 6G–J and S3 Fig.). In addition, RAB11A-siRNA-treated cells exhibited cytokinesis defects, such as formation of binucleated cells and too few membrane blebs (data not shown).

**Figure 6.**
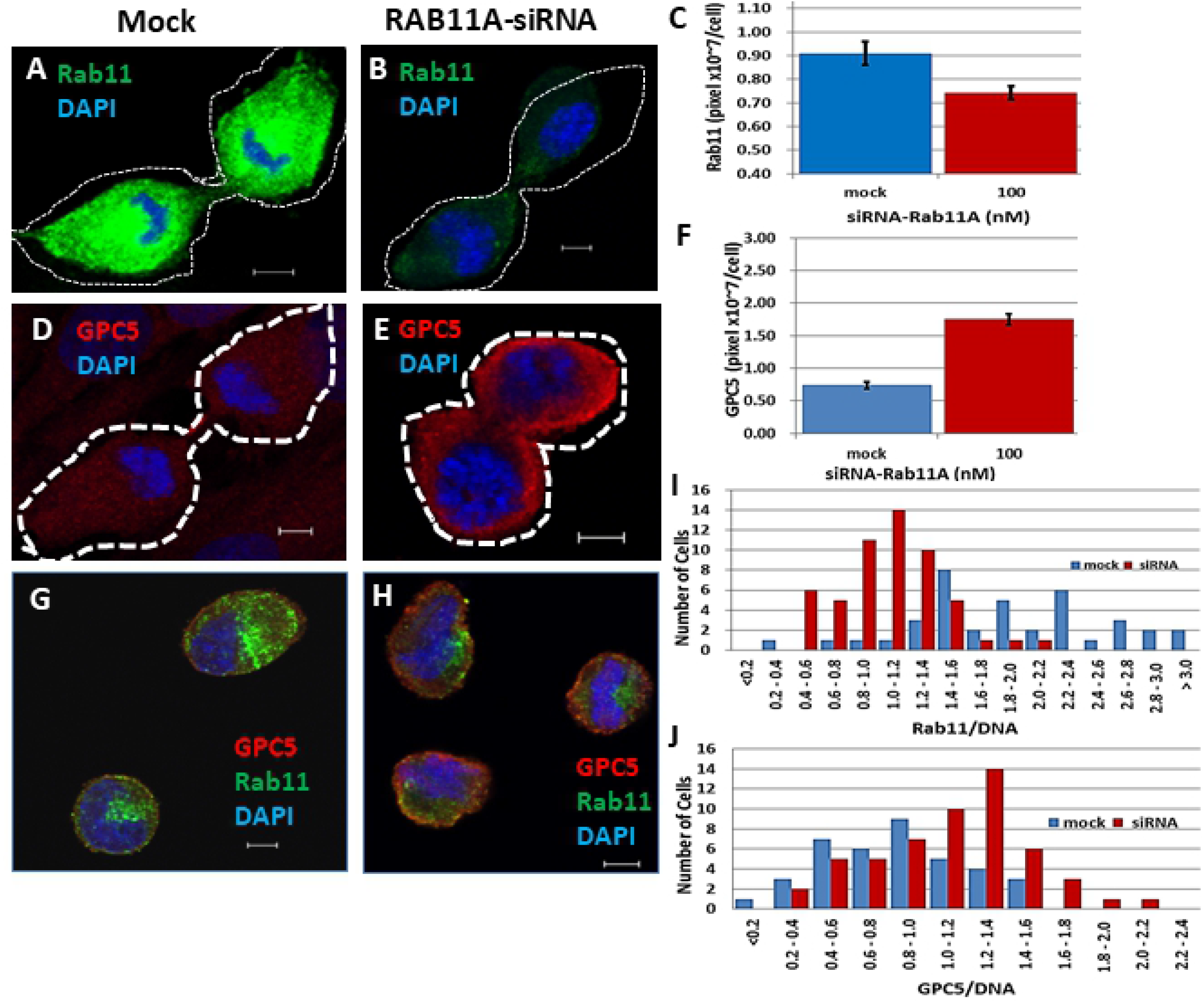
Localization of GPC5 and Rab11 in U3DT cells treated with RAB11A-siRNA. (A–B) Images of Rab11 in untreated (A) and RAB11A-siRNA-treated U3DT cells (B). (C) Quantitative analysis of U3DT cells treated (red) or not treated (blue) with 100 nM RAB11A-siRNA. (D–E) Images of GPC5 in U3DT cells not treated (D) or treated (E) with 100 nM RAB11A-siRNA (E). (F) Quantitative expression of GPC5 in U3DT cells not treated (blue) or treated (red) with 100 nM RAB11A-siRNA. (G–H) Images of GPC5 (red) and Rab11 (green) in U3DT cells not treated (G) or treated (H) with RAB11A-siRNA. (I–J) Immunofluorescence images of trypsinized cells were obtained using a Leica SP-8 immunofluorescence microscope, and the pixel sum was estimated with the SP8 software (RAS X). A histogram of Rab11 images is shown for each untreated (blue) and RAB11A-siRNA-treated U3DT cell (red) (I). A histogram of GPC5 images is shown for each untreated (blue) and RAB11A-siRNA-treated U3DT cell (red) (J). Scale bar, 5 μm.

### GPC5 is released into the conditioned medium

Circulating exosomes positive for GPC1 have been isolated from the serum of patients with pancreatic ductal adenocarcinoma cells [31]. Therefore, we assessed whether GPC5 could be detected in the conditioned medium of U3DT cells. We isolated EVs from the conditioned medium of U3DT cells by differential ultracentrifugation and filtration. The EV population was mixed, and electron microscopy revealed a size distribution with a peak diameter of ca. 100 nm (Fig. 7A–B). EVs immunostained with anti-Rab11 (Fig. 7C), anti-FGFR (Fig. 7F), anti-CD63 (Fig. 7I), and ant-ARF6 (Fig. 7L) antibodies exhibited significant colocalizations with GPC5, as supported by line scan determination (Fig. 7D, G, J, and M). EVs colocalized with Rab11 or FGFR in GPC5-containing EVs were 94% (Fig. 7E and H), whereas EVs with CD63 or ARF6 in them were 25% and 44% (Fig. 7K and N), respectively (S4 Fig.). These results indicate that GPC5-containing EVs generally contain Rab11 or FGFR, but do not include so many CD63 or ARF6.

**Figure 7.**
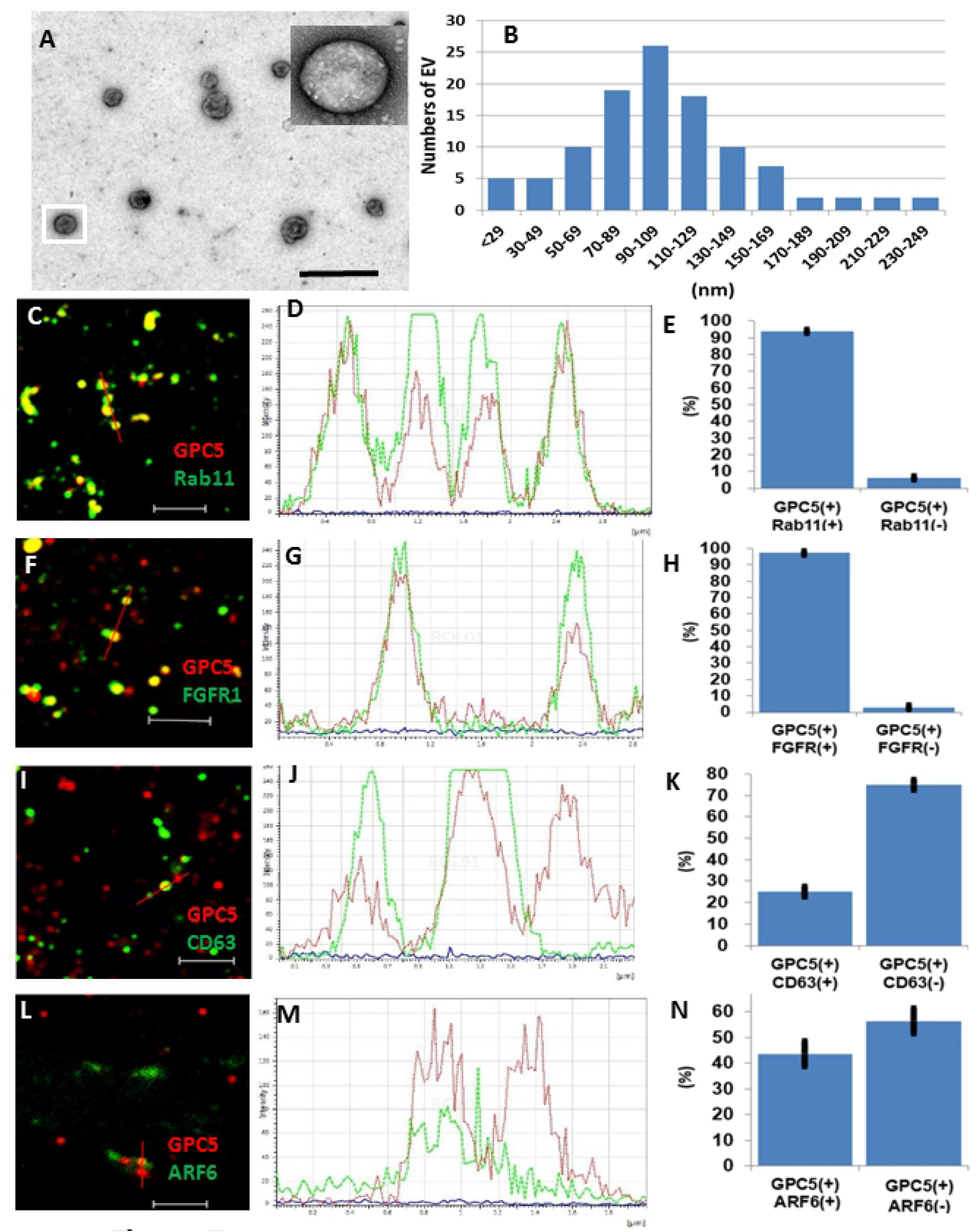
GPC5 is present on EVs from U3DT cells. (A) Electron microscopic image of negative-stained EVs. The inset shows a magnification of the boxed EV. (B) Size distribution of vesicles in EV preparations measured by image software (n = 110). (C) (F) (I) (L) GPC5-immunostained EVs (red) stained for Rab11 (green) (C), FGFR1 (green) (F), CD63 (green) (I), and ARF6 (green) (L). (D) (G) (J) (M) Line scan determination of the red bars in (C), (F), (I), and (L): GPC5 (red), others (green). (E) (H) (K) (N) Quantification of scan determinations in (D), (G), (J), and (M). n = 1,516, (E), 630 (H), 704 (K), and 406 (N). Scale bars: (A), 500 nm; (C), (F), (I), (L), 2 μm.

Previously, diverse biological functions have been attributed to EVs [32]. To examine the functional interactions of EVs with cells, we tested U3DT-EVs for their ability to deliver GPC5 to UE6E7T-3 cells, the parental cell line of U3DT, which expresses very little if any GPC5. After culture of UE6E7T-3 cells with U3DT-EVs for 25 h, cells double-stained with anti-GPC5 and anti-FGFR1 antibodies were positive for GPC5 in the perinuclear region (Fig. 8C), and the GPC5 signal overlapped with the FGFR1 signal in this region (Fig. 8D). Flow-cytometric analysis also revealed extensive uptake of the U3DT-EVs by UE6E7T-3 cells (Fig. 8F–G).

**Figure 8.**
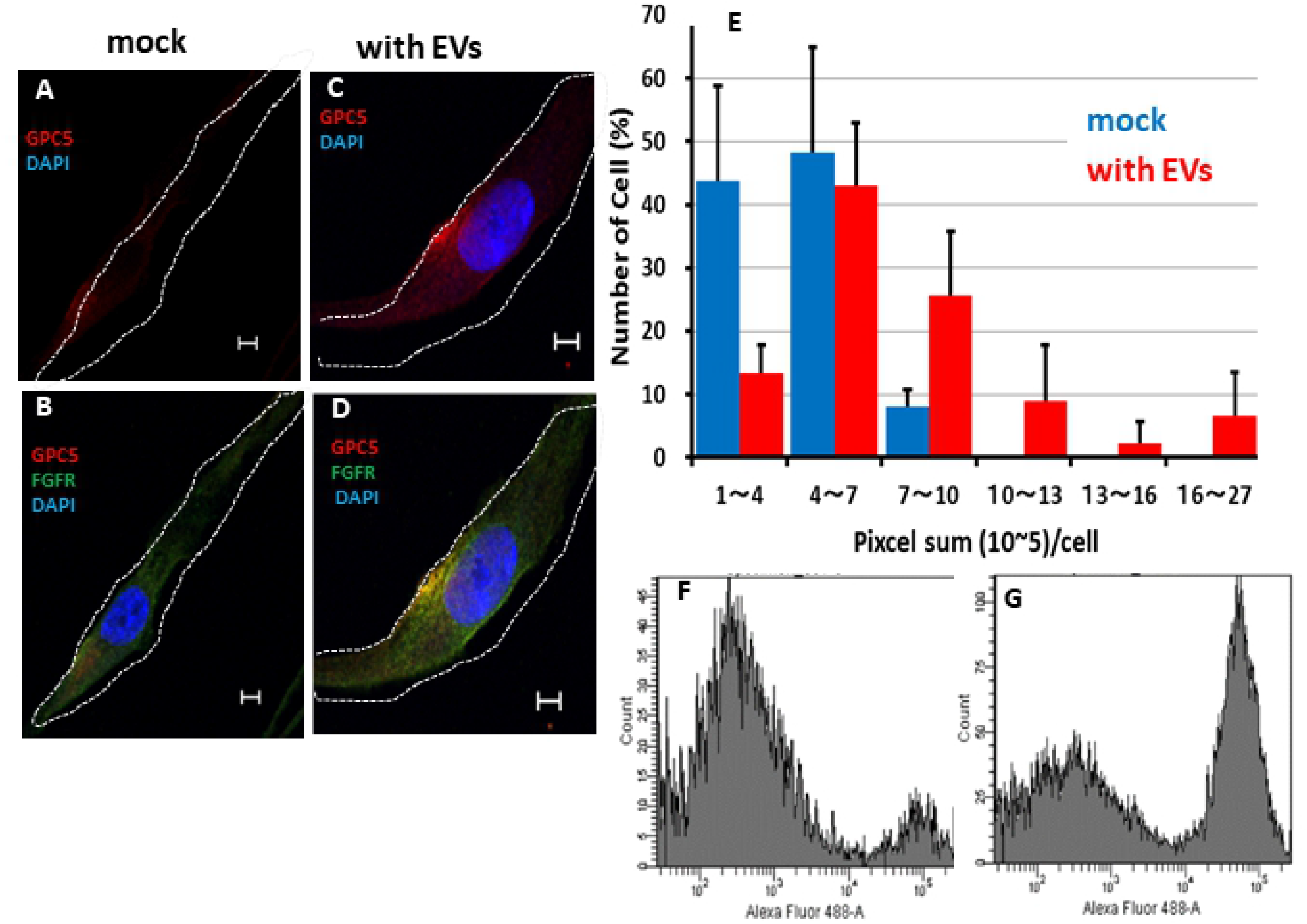
EVs are incorporated to UE6E7T3 cells. (A–B) Images of control cells. (C–D) Images of EV-treated cells. (B) (D) Merged images of UE6E7T-3 cells (B) and EV-treated cells (D) stained for GPC5 (red) and FGFR1 (green). (E) Histogram of GPC5 (pixel sum per cell) in control UE6E7T-3 cells (blue) or of cells cultured with EVs for 1 day (red). n = 29 (blue) and 46 (red). (F–G) FACS pattern of GPC5 in UE6E7T-3 cells (F) and in cells cultured with EVs for 1 day (G). Scale bar, 5 μm.

## Discussion

The results of this study reveal novel subcellular localizations of GPC5 in association with dynamic cell motility and cell communication. Although understanding the detailed functions of GPC5 will require further studies at the molecular level, we clearly demonstrated that GPC5 contributes to cell motility and cell communication. Our recent gene expression analysis of U3DT cells revealed that GPC5 colocalizes with Ptc1 in the perinuclear region. In this study of the subcellular distribution of GPC5, we found that the protein markedly accumulated at the leading edge of cell migration, intercellular bridge, and bleb protrusions during cytokinesis, as well as in EVs released into the conditioned medium (Fig. 9).

**Figure 9.**
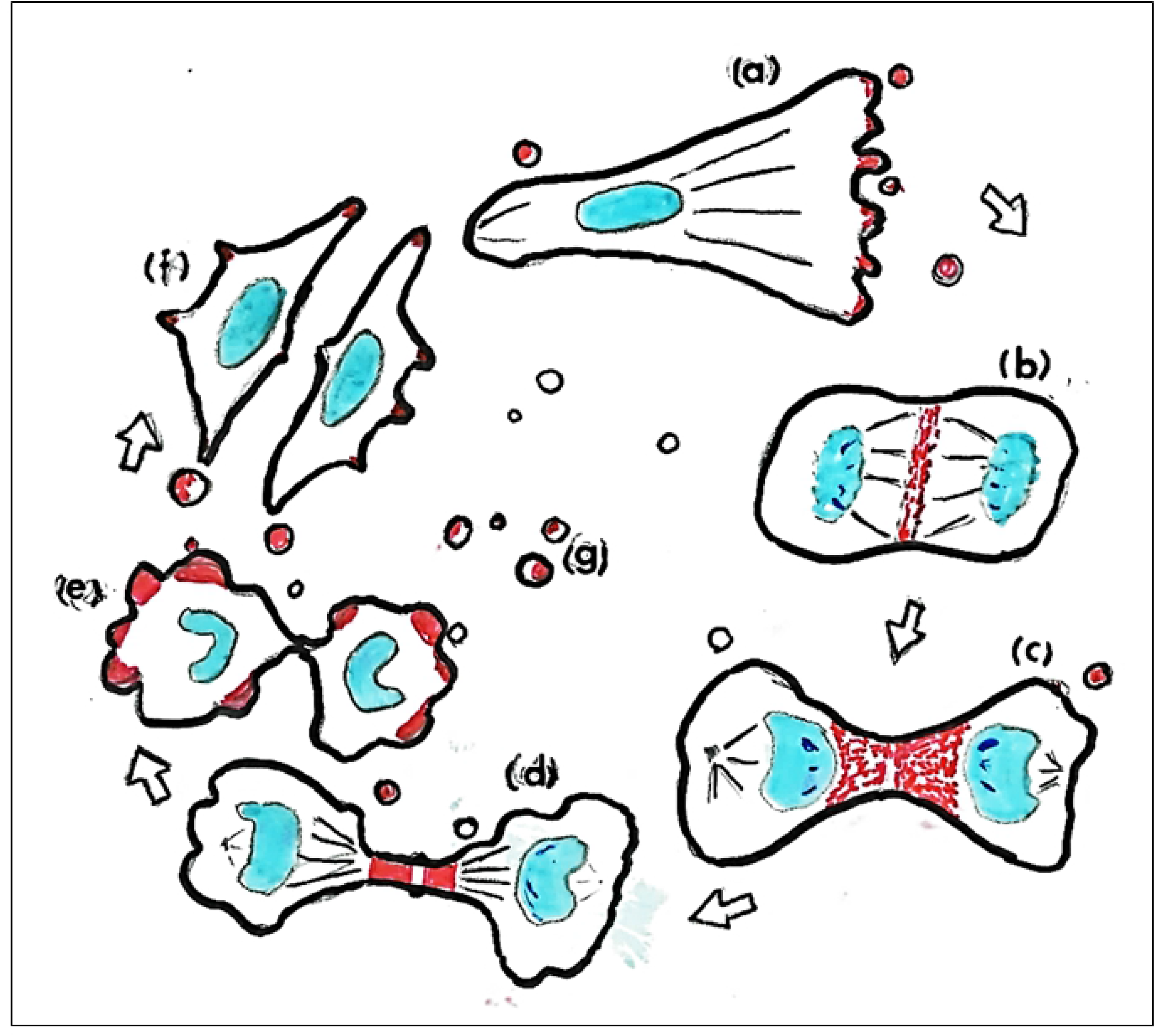
Summary. Subcellular localization of GPC5 during the cell cycle. (A) GPC5 (red) localizes at the leading edge of cell migration, (B) at the equatorial plane, (C) at the furrow, (D) at the intercellular bridge during mitosis, (E) in membrane blebs during cytokinesis, (F) at the tips of filopodia, and (G) in EVs.

Cell-surface HSPGs contribute to signal transduction as co-receptors, as well as to the stable retention of multiple growth factors. To date, the contribution of HSPGs to cell migration has mostly been studied in regard to the involvement of SDC4 in FGF-induced migration. SDC4 mediates the matrix-induced protein kinase Cα(PKCα)-dependent activation of Rac1 (Rho family of GTPase) and localizes Rac1 to the leading edge of migrating cells, thereby promoting migration [33]. However, SDC4-induced cell migration can occur even in the presence of dominant-negative FGFR1 [34]. In this study, we demonstrated strong colocalization of GPC5 and FGFR1 at the leading edges of migrating cells in response to unknown stimuli. This phenomenon may be induced by the FGF-stimulated FGFR1 signaling pathway, in which activated FGFR1 stimulates P13-kinase, resulting in Rac1 activation and leading in turn to migration [35]. The pathway is different from the SDC4-induced PKCα-dependent signaling pathway [36]. Thus, we is speculate that GPC5 promotes the migration of U3DT cells via the FGFR1 signaling pathway.

Further evidence that GPC5 also colocalizes with Rab11 or ARF6 at the leading edge of cell migration suggest recycle endosomal dynamics in U3DT cells (Fig. 2). Both Rab11 and ARF6 are well known for their roles in regulating vesicular transport and cytoskeletal dynamics [37-39]. In cultured human cells, Rab11 transports membrane cargos to and from recycling endosomes on the way to the plasma membrane [20,40], whereas ARF6 induces the formation of large lamellipodia and promotes cell migration [41], suggesting the involvement of the Rab11 or ARF6 recycling pathways in the transport of GPC5 and FGFR1 in U3DT cell migration.

Interestingly, we found that GPC5 was enriched in the intercellular bridge connecting the daughters during division of U3DT cells. During cytokinesis, many factors necessary for abscission localize to the cleavage furrow and midbody through a variety of pathways, including endosomal recycling. The Rab11-dependent recycling pathways influence a wide range of cell-surface proteins. Because Rab11-positive vesicles separate from Rab11-containing recycling endosomes and traffic to the cleavage furrow and midbody, we first sought to determine whether GPC5 associates with Rab11. Three lines of evidence suggest this possibility. First, GPC5 colocalizes with Rab11 at intercellular bridges in late telophase. GPC5 is probably transported to the intercellular bridge in association with Rab11-endosomes. Second, depletion of Rab11 using RAB11A-siRNA gave rise to accumulation of GPC5 inside the cell, as demonstrated by immunostaining. This is consistent with previous studies showing that Rab11 depletion inhibits exocytic events of recycling vesicles [20,42,43]. Interestingly, GPC5-containing vesicle-like dots that have yet to start budding were also observed beneath the cell surface (Fig. 7H and S2 Fig.). This image seems to reveal an intermediate process in the budding pathway, suggesting that the budded vesicles are generated from GPC5-Rab11-containing dots. Third, vesicles containing GPC5 and Rab11, ranging from 40 to 250 nm in diameter, were released from U3DT cells. Taken together, our data strongly suggest that cell-surface GPC5 is transported in association with Rab11 inside cells.

Various organelles and components participate in cytokinesis, but it remains to be elucidated how each component (or organelle) interacts with the others. Although we demonstrated that GPC5 is located at the intercellular bridge during cell division, we could not elucidate the role of GPC5 during cytokinesis. Dynamic blebbing was observed during cytokinesis in U3DT cells. Blebbing appeared at late anaphase and disappeared after cytokinesis was completed, and the two cells entered interphase. GPC5 accumulated in blebs, and in some blebs the signal was more condensed at the leading edge. This may indicate that GPC5 is correlated with the initial forward movement in blebs, as observed at the leading edge of cell migration. The role of blebbing during cytokinesis is unknown. One possible explanation is that polarized blebbing movement induces mechanical separation of the daughter cells [44]. Support for this speculation comes from evidence that blebs tend to form at poles, away from the cleavage furrow (S1 Fig.). Blebbing is a dynamic and rapid amoeboid movement, and faster-separating cells exert a stronger pull on their intercellular bridge [45].

Bleb growth decreased upon treatment of U3DT cells with RAB11A-siRNA. One possible explanation for this is that membrane sources for blebbing are diminished upon Rab11 inhibition. Bleb inflation is the result of unfolding of stored membrane, which could originate from fusion of Rab11 vesicles with plasma membrane [46,47]. Alternatively, Rab11 inhibition may prevent cytoplasmic flow from pushing the membrane outwards.

GPC1-containing endosomes in the serum are a potential marker for early pancreatic cancer [31]. In this study, we detected EVs containing GPC5 in the conditioned medium of U3DT cells. These EVs were a heterogeneous mixture, varying in size from 40 to 250 nm. Moreover, we confirmed by immunostaining that GPC5-containing EVs associate with the exosome/microvesicle markers Rab11 and ARF6. Rab11 is required for exosome secretion, microvesicle budding, and viral budding [42,43]. However, our results show that Rab11 not only promotes EV-budding, but is budded along with GPC5 into the EVs themselves. How GPC5 and Rab11 are recruited into EVs remains unknown.

Accumulating evidence indicate that EVs generated by highly aggressive cancer cells are capable of promoting tumor growth [47-49]. We showed in Fig. 8 that GPC5-containing EVs of U3DT cells are taken in UE6E7T-3 cells, which do not express detectable GPC5, as indicated in Fig. 1. Previous studies by our group and others have shown that GPC5 significantly stimulate cell proliferation. However, some GPC5-containing EV preparations had an increased proliferative effect on UE6E7T-3 cells, whereas other preparations had no effect (data not shown). It is likely that the heterogeneity of EVs is one of the main causes for this. Further work with GPC5-containing EVs is needed to confirm the effects of these vesicles on cell growth.

Previously, we showed that U3DT cells transformed after prolonged culture express significantly elevated levels of GPC5, localized to the plasma membrane. In this study, we showed that GPC5 also accumulates at the leading edges of migrating cells, intercellular bridges, and membrane blebs during cell division, and that EVs are released from U3DT cells. Although the subcellular localization of GPC5 might be crucial for its functions, each function in turn might also play important role in promoting tumorigenesis by U3DT cells. Williamson et al. showed that GPC5 stimulates the proliferation of RMS cells [4]. Thus, the evidence obtained in this study should facilitate assessment of the functional contribution of GPC5-promoting proliferation in sarcoma, as well as the usefulness of EV as a biomarker for sarcoma.

## Conclusions

GPC5 localized not only at primary cilia on cell surface, but also at the leading edge of migrating cells, at the intercellular bridge and blebs during cytokinesis, and in EVs. The evidences might provide hints in assessment of the functional contribution of GPC5-promoting proliferation [Fig. 9].

## Acknowledgements

We thank H. Kamada, T.S. Kodama and Y. Kasahara of the National Institute of Biomedical Innovation for encouraging our research during this study. We are grateful to FUJIFILM Wako Pure Chemical Co. and Santa Cruz Biotechnology for donation of many sample antibodies.

## Supporting information

**S1 Fig. Many blebs are observed outside two daughter cells.** U3DT cell at telophase was stained with anti-GPC5 antibody (red), Alexa Fluor 488-WGA (green), anti-Rab11A antibodies (grey) and DAPI (blue). The immunostained U3DT cells were observed with Leica SP8 immunofluorescence microscope and their images were obtained with sectioning of 0.235μm (A) or 0.362μm (B). Scale bar, 5 μm.

**S2 Fig. Blebs containing GPC5 and Rab11 are observed at telophase.** U3DT cell at telophase was stained with anti-GPC5 antibody (red), Alexa Fluor 488-WGA (green), anti-Rab11A antibodies (grey) and DAPI (blue). Scale bar, 2 μm.

**S3 Fig. Localization of GPC5 increases on surface of U3DT cells treated with RAB11A-siRNA.** Immunofluorescence images of trypsinized cells were obtained using a Leica SP-8 immunofluorescence microscope. Images of GPC5 (red), and Rab11 (green) and DAPI (blue) in U3DT cells not treated (left) or treated (right) with RAB11A-siRNA. Scale bar, 5 μm.

**S4 Fig. EVs contain GPC5, FGFR1 and Rab11.** Expand images of Fig. 7. (C) (F) (I) (L): GPC5-immunostained EVs (red) stained for Rab11 (green) (C), FGFR1 (green) (F), CD63 (green) (I), and ARF6 (green) (L). (D) (G) (J) (M) Line scan determination of the red bars in (C), (F), (I), and (L): GPC5 (red), others (green). Scale bars: 2 μm.

## AUTHOR CONTRIBUTIONS

MT, KT, J-KT and KA contributed to the design of this research. MT, KT, YM and J-KT performed the experiments. MT, KT, YM, TT, RY, J-KT and KA contributed to data analysis. MT and KT wrote the manuscript. MT, KT, J-KT and KA contributed equally to this work. All authors read and approved the final manuscript.

## References

1. Sarrazin S, Lamanna WC, Esko JD. Heparan sulfate proteoglycans. Cold Spring Harbor Perspectives in Biology. 2011;3: 1–33. doi:10.1101/cshperspect.a004952

2. Li N, Gao W, Zhang YF, Ho M. Glypicans as Cancer Therapeutic Targets. Trends in Cancer. 2018;4: 741–754. doi:10.1016/j.trecan.2018.09.004

3. Takeuchi M, Higashino A, Takeuchi K, Hori Y, Koshiba-Takeuchi K, Makino H, et al. Transcriptional dynamics of immortalized human mesenchymal stem cells during transformation. PLoS ONE. 2015;10: 1–23. doi:10.1371/journal.pone.0126562

4. Williamson D, Selfe J, Gordon T, Lu YJ, Pritchard-Jones K, Murai K, et al. Role for amplification and expression of Glypican-5 in rhabdomyosarcoma. Cancer Research. 2007;67: 57–65. doi:10.1158/0008-5472.CAN-06-1650

5. Li F, Shi W, Capurro M, Filmus J. Glypican-5 stimulates rhabdomyosarcoma cell proliferation by activating Hedgehog signaling. Journal of Cell Biology. 2011;192: 691–704. doi:10.1083/jcb.201008087

6. Witt RM, Hecht ML, Pazyra-Murphy MF, Cohen SM, Noti C, Van Kuppevelt TH, et al. Heparan sulfate proteoglycans containing a glypican 5 core and 2-O-sulfo-iduronic acid function as sonic hedgehog co-receptors to promote proliferation. Journal of Biological Chemistry. 2013;288: 26275–26288. doi:10.1074/jbc.M112.438937

7. Zhang Y, Wang J, Dong F, Li H, Hou Y. The role of GPC5 in lung metastasis of salivary adenoid cystic carcinoma. Archives of Oral Biology. 2014;59: 1172–1182. doi:10.1016/j.archoralbio.2014.07.009

8. Sun Y, Xu K, He M, Fan G, Lu H. Overexpression of glypican 5 (GPC5) inhibits prostate cancer cell proliferation and invasion via suppressing Sp1-mediated EMT and activation of Wnt/β-catenin signaling. Oncology Research. 2018;26: 565–572. doi:10.3727/096504017X15044461944385

9. Yuan S, Yu Z, Liu Q, Zhang M, Xiang Y, Wu N, et al. GPC5, a novel epigenetically silenced tumor suppressor, inhibits tumor growth by suppressing Wnt/β-catenin signaling in lung adenocarcinoma. Oncogene. 2016;35: 6120–6131. doi:10.1038/onc.2016.149

10. Li Y, Miao L, Cai H, Ding J, Xiao Y, Yang J, et al. The overexpression of glypican-5 promotes cancer cell migration and is associated with shorter overall survival in non-small cell lung cancer. Oncology Letters. 2013;6: 1565–1572. doi:10.3892/ol.2013.1622

11. Yang X, Zhang Z, Qiu M, Hu J, Fan X, Wang J, et al. Glypican-5 is a novel metastasis suppressor gene in non-small cell lung cancer. Cancer Letters. 2013;341: 265–273. doi:10.1016/j.canlet.2013.08.020

12. Guo L, Wang J, Zhang T, Yang Y. Glypican-5 is a tumor suppressor in non-small cell lung cancer cells. Biochemistry and Biophysics Reports. 2016;6: 108–112. doi:10.1016/j.bbrep.2016.03.010

13. Ornitz DM, Itoh N. The fibroblast growth factor signaling pathway. Wiley Interdisciplinary Reviews: Developmental Biology. 2015;4: 215–266. doi:10.1002/wdev.176

14. Kleeff J, Ishiwata T, Kumbasar A, Friess H, Büchler MW, Lander AD, et al. The cell-surface heparan sulfate proteoglycan glypican-1 regulates growth factor action in pancreatic carcinoma cells and is overexpressed in human pancreatic cancer. Journal of Clinical Investigation. 1998;102: 1662–1673. doi:10.1172/JCI4105

15. Huang G, Ge G, Izzi V, Greenspan DS. α3 Chains of type v collagen regulate breast tumour growth via glypican-1. Nature Communications. 2017;8: 1–17. doi:10.1038/ncomms14351

16. Su G, Meyer K, Nandini CD, Qiao D, Salamat S, Friedl A. Glypican-1 is frequently overexpressed in human gliomas and enhances FGF-2 signaling in glioma cells. American Journal of Pathology. 2006;168: 2014–2026. doi:10.2353/ajpath.2006.050800

17. Tkachenko E, Lutgens E, Stan RV, Simons M. Fibroblast growth factor 2 endocytosis endothelial cells proceed via syndecan-4-dependent activation of Rac1 and a Cdc42-dependent macropinocytic pathway. Journal of Cell Science. 2004;117: 3189–3199. doi:10.1242/jcs.01190

18. Couchman JR, Multhaupt H, Sanderson RD. Recent insights into cell surface heparan sulphate proteoglycans and cancer [version 1; referees: 3 approved]. F1000Research. 2016;5: 1–9. doi:10.12688/F1000RESEARCH.8543.1

19. Kanada M, Bachmann MH, Hardy JW, Frimannson DO, Bronsart L, Wang A, et al. Differential fates of biomolecules delivered to target cells via extracellular vesicles. Proceedings of the National Academy of Sciences of the United States of America. 2015;112: E1433–E1442. doi:10.1073/pnas.1418401112

20. Takahashi S, Kubo K, Waguri S, Yabashi A, Shin HW, Katoh Y, et al. Rab11 regulates exocytosis of recycling vesicles at the plasma membrane. Journal of Cell Science. 2012;125: 4049–4057. doi:10.1242/jcs.102913

21. Yu B, Yang H, Zhang X, Li H. Visualizing and quantifying the effect of the inhibition of HSP70 on breast cancer cells based on laser scanning microscopy. Technology in Cancer Research and Treatment. 2018;17: 1–7. doi:10.1177/1533033818785274

22. Rieck PW, Cholidis S, Hartmann C. Intracellular signaling pathway of FGF-2-modulated corneal endothelial cell migration during wound healing in vitro. Experimental Eye Research. 2001;73: 639–650. doi:10.1006/exer.2001.1067

23. Nikolovska K, Spillmann D, Seidler DG. Uronyl 2-O sulfotransferase potentiates Fgf2-induced cell migration. Journal of Cell Science. 2015;128: 460–471. doi:10.1242/jcs.152660

24. Volk R, Schwartz JJ, Li J, Rosenberg RD, Simons M. The role of syndecan cytoplasmic domain in basic fibroblast growth factor-dependent signal transduction. Journal of Biological Chemistry. 1999;274: 24417–24424. doi:10.1074/jbc.274.34.24417

25. Donaldson JG, Porat-Shliom N, Cohen LA. Clathrin-independent endocytosis: A unique platform for cell signaling and PM remodeling. Cellular Signalling. 2009;21: 1–6. doi:10.1016/j.cellsig.2008.06.020

26. Chen TC, Lee SA, Hong TM, Shih JY, Lai JM, Chiou HY, et al. From midbody protein - Protein interaction network construction to novel regulators in cytokinesis. Journal of Proteome Research. 2009;8: 4943–4953. doi:10.1021/pr900325f

27. Horgan CP, Walsh M, Zurawski TH, McCaffrey MW. Rab11-FIP3 localises to a Rab11-positive pericentrosomal compartment during interphase and to the cleavage furrow during cytokinesis. Biochemical and Biophysical Research Communications. 2004;319: 83–94. doi:10.1016/j.bbrc.2004.04.157

28. Wilson GM, Fielding AB, Simon GC, Yu X, Andrews PD, Haines RS, et al. The FIP3-Rab11 protein complex regulates recycling endosome targeting to the cleavage furrow during late cytokinesis. Molecular Biology of the Cell. 2005;16: 849–860. doi:10.1091/mbc.E04-10-0927

29. Yoshigaki T. Accumulation of WGA receptors in the cleavage furrow during cytokinesis of sea urchin eggs. Experimental Cell Research. 1997;236: 463–471. doi:10.1006/excr.1997.3740

30. Boucrot E, Kirchhausen T. Endosomal recycling controls plasma membrane area during mitosis. Proceedings of the National Academy of Sciences of the United States of America. 2007;104: 7939–7944. doi:10.1073/pnas.0702511104

31. Melo SA, Luecke LB, Kahlert C, Fernandez AF, Gammon ST, Kaye J, et al. Glypican-1 identifies cancer exosomes and detects early pancreatic cancer. Nature. 2015;523: 177–182. doi:10.1038/nature14581

32. Maas SLN, Breakefield XO, Weaver AM. Extracellular Vesicles: Unique Intercellular Delivery Vehicles. Trends in Cell Biology. 2017;27: 172–188. doi:10.1016/j.tcb.2016.11.003

33. Bass MD, Roach KA, Morgan MR, Mostafavi-Pour Z, Schoen T, Muramatsu T, et al. Syndecan-4-dependent Rac1 regulation determines directional migration in response to the extracellular matrix. Journal of Cell Biology. 2007;177: 527–538. doi:10.1083/jcb.200610076

34. Tkachenko E, Elfenbein A, Tirziu D, Simons M. Syndecan-4 clustering induces cell migration in a PDZ-dependent manner. Circulation Research. 2006;98: 1398–1404. doi:10.1161/01.RES.0000225283.71490.5a

35. Kanazawa S, Fujiwara T, Matsuzaki S, Shingaki K, Taniguchi M, Miyata S, et al. bFGF regulates PI3-Kinase-Rac1-JNK pathway and promotes fibroblast migration in wound healing. PLoS ONE. 2010;5: 1–12. doi:10.1371/journal.pone.0012228

36. Horowitz A, Tkachenko E, Simons M. Fibroblast growth factor-specific modulation of cellular response by syndecan-4. Journal of Cell Biology. 2002;157: 715–725. doi:10.1083/jcb.200112145

37. Fielding AB, Schonteich E, Matheson J, Wilson G, Yu X, Hickson GRX, et al. Rab11-FIP3 and FIP4 interact with Arf6 and the Exocyst to control membrane traffic in cytokinesis. EMBO Journal. 2005;24: 3389–3399. doi:10.1038/sj.emboj.7600803

38. Jing J, Tarbutton E, Wilson G, Prekeris R. Rab11-FIP3 is a Rab11-binding protein that regulates breast cancer cell motility by modulating the actin cytoskeleton. European Journal of Cell Biology. 2009;88: 325–341. doi:10.1016/j.ejcb.2009.02.186

39. Welz T, Wellbourne-Wood J, Kerkhoff E. Orchestration of cell surface proteins by Rab11. Trends in Cell Biology. 2014;24: 407–415. doi:10.1016/j.tcb.2014.02.004

40. Sato M, Grant BD, Harada A, Sato K. Rab11 is required for synchronous secretion of chondroitin proteoglycans after fertilization in Caenorhabditis elegans. Journal of Cell Science. 2008;121: 3177–3186. doi:10.1242/jcs.034678

41. Santy LC, Casanova JE. Activation of ARF6 by ARNO stimulates epithelial cell migration through downstream activation of both Rac1 and phospholipase D. Journal of Cell Biology. 2001;154: 599–610. doi:10.1083/jcb.200104019

42. Bruce EA, Digard P, Stuart AD. The Rab11 Pathway Is Required for Influenza A Virus Budding and Filament Formation. Journal of Virology. 2010;84: 5848–5859. doi:10.1128/jvi.00307-10

43. Wehman AM, Poggioli C, Schweinsberg P, Grant BD, Nance J. The P4-ATPase TAT-5 inhibits the budding of extracellular vesicles in C. elegans embryos. Current Biology. 2011;21: 1951–1959. doi:10.1016/j.cub.2011.10.040

44. Sedzinski J, Biro M, Oswald A, Tinevez JY, Salbreux G, Paluch E. Polar actomyosin contractility destabilizes the position of the cytokinetic furrow. Nature. 2011;476: 462–468. doi:10.1038/nature10286

45. Lafaurie-Janvore J, Maiuri P, Wang I, Pinot M, Manneville J-B, Betz T, et al. ESCRT-III Assembly and Cytokinetic Abscission Are Induced by Tension Release in the Intercellular Bridge. Science. 2013;339: 1625 LP–1629. doi:10.1126/science.1233866

46. Ai E, Skop AR. Endosomal recycling regulation during cytokinesis. Communicative and Integrative Biology. 2009;2: 444–447. doi:10.4161/cib.2.5.8931

47. Goudarzi M, Tarbashevich K, Mildner K, Begemann I, Garcia J, Paksa A, et al. Bleb Expansion in Migrating Cells Depends on Supply of Membrane from Cell Surface Invaginations. Developmental Cell. 2017;43: 577–587.e5. doi:10.1016/j.devcel.2017.10.030

48. Heneberg P. Paracrine tumor signaling induces transdifferentiation of surrounding fibroblasts. Critical Reviews in Oncology/Hematology. 2016;97: 303–311. doi:10.1016/J.CRITREVONC.2015.09.008

49. Choi D, Lee TH, Spinelli C, Chennakrishnaiah S, D’Asti E, Rak J. Extracellular vesicle communication pathways as regulatory targets of oncogenic transformation. Seminars in Cell and Developmental Biology. 2017;67: 11–22. doi:10.1016/j.semcdb.2017.01.003

